# Ivermectin reduces coronavirus infection *in vivo*: a mouse experimental model

**DOI:** 10.1101/2020.11.02.363242

**Authors:** AP Arévalo, R Pagotto, J Pórfido, H Daghero, M Segovia, K Yamasaki, B Varela, M Hill, JM Verdes, M Duhalde Vega, M Bollati-Fogolín, M Crispo

**Affiliations:** Transgenic and Experimental Animal Unit, Institut Pasteur de Montevideo, Uruguay; Cell Biology Unit, Institut Pasteur de Montevideo, Uruguay; Laboratory of Immunoregulation and Inflammation, Institut Pasteur de Montevideo, Uruguay; Pathobiology Department, Faculty of Veterinary, Montevideo, Uruguay; Institute of Biological Chemistry and Chemical Physics (UBA-CONICET). School of Pharmacy and Biochemistry, University of Buenos Aires, Argentina

**Keywords:** SARS-CoV2, MHV, IVM, liver, mice.

## Abstract

SARS-CoV2 is a single strand RNA virus member of the type 2 coronavirus family, responsible for causing COVID-19 disease in humans. The objective of this study was to test the ivermectin drug in a murine model of coronavirus infection using a type 2 family RNA coronavirus similar to SARS-CoV2, the mouse hepatitis virus (MHV). BALB/cJ female mice were infected with 6,000 PFU of MHV-A59 (Group Infected; n=20) and immediately treated with one single dose of 500 μg/kg of ivermectin (Group Infected + IVM; n=20), or were not infected and treated with PBS (Control group; n=16). Five days after infection/treatment, mice were euthanized to obtain different tissues to check general health status and infection levels. Overall results demonstrated that viral infection induces the typical MHV disease in infected animals, with livers showing severe hepatocellular necrosis surrounded by a severe lymphoplasmacytic inflammatory infiltration associated with a high hepatic viral load (52,158 AU), while ivermectin administration showed a better health status with lower viral load (23,192 AU; p<0.05) and few livers with histopathological damage (p<0.05), not showing statistical differences with control mice (P=NS). Furthermore, serum transaminase levels (aspartate aminotransferase and alanine aminotransferase) were significantly lower in treated mice compared to infected animals. In conclusion, ivermectin seems to be effective to diminish MHV viral load and disease in mice, being a useful model for further understanding new therapies against coronavirus diseases.

## Introduction

SARS-CoV2 is a beta-coronavirus recently identified to be responsible for the severe acute respiratory syndrome COVID-19. This new pandemic virus is causing huge losses in terms of human lives and macroeconomic consequences worldwide. Tremendous efforts are being carried out to stop the pandemic that is affecting nowadays millions of people, and in that sense early and accurate diagnosis, effective vaccines or proven treatments are being developed. However, experimental conditions with SARS-CoV2 are not easily available in research laboratories due to biosecurity reasons, thus having *in vivo* preclinical data is not always easy. Mouse hepatitis virus (MHV) is a similar single strand RNA coronavirus (Fan et al., 2020) affecting different mouse organs (Weiss and Leibowitz, 2011), also highly contagious with natural transmission occurring by respiratory or oral routes, showing high morbidity and low mortality rate, with no vaccine or treatment available, whose control requires to sacrifice the entire laboratory mice colony when an infection occurs. It has been proposed that MHV could be an interesting model to test new therapies for COVID 19 in animal models, since it has been recently demonstrated that the mechanism of infection has some similarities with SARS-CoV-2 (Körner et al., 2020). After entry into the host cell, both coronaviruses require a similar nuclear transport system mediated by the importin α/β1 heterodimer (Timani et al., 2005; Wulan et al., 2015), making this system an interesting target for the development of candidate therapies against the viral infection.

Ivermectin is an efficient and non-expensive drug usually applied to treat parasite infestations, FDA-approved for animal and human use and available worldwide. It has been proved to have a wide margin of safety with a DL_50_ of 30 mg/kg in mice and is used in humans at a therapeutic dose of 150-200 μg/kg as antiparasitic treatment (Crump and Ōmura, 2011). This drug acts on the cells at different levels, and in some cases has shown an *in vitro* effect against RNA and DNA virus infection (Heidary and Gharebaghi, 2020) by the suppression of a host cellular process related with the inhibition of nuclear transport of specific proteins required for viral replication (Wagstaff et al., 2012). Recently, in June 2020, it was reported in an *in vitro* cell model that ivermectin was effective against SARS-CoV2, showing an inhibition of the virus replication and making it a possible candidate for COVID-19 as a repurposing drug (Caly et al., 2020). Information on the *in vivo* antiviral effect of ivermectin against coronavirus has not been published yet, something needed in order to progress on the development of new therapeutic strategies for the control of these types of coronavirus.

The objective of this study was to evaluate the *in vivo* effect of the ivermectin drug in a murine model of a type 2 family RNA coronavirus, the MHV, in terms of general health profile and hepatic viral load and functionality. We hypothesize that the administration of a single dose of ivermectin in recently infected mice decreases viral load and impairs the action of the virus on the host organism.

## Materials and Methods

### Animals and management

A total of 56 BALB/cJ female mice (6-8 weeks old) were bred at the Transgenic and Experimental Animal Unit of Institut Pasteur de Montevideo, under specific pathogen free conditions in individually ventilated racks (IVC, 1285L, Tecniplast, Milan, Italy). During the experimental procedure, females were housed in groups of seven in negative pressure microisolators (ISOCageN, Tecniplast) with aspen wood bedding chips (Toplit 6, SAFE, Augy, France), paper towels and cardboard tubes as environmental enrichment. They had *ad libitum* access to autoclaved food (5K67, LabDiet, MO, USA) and filtered water. Housing environmental conditions during the experiment were as follow: 20±1°C temperature, 30-70% relative humidity, negative pressure (biocontainment) and light/dark cycle of 12/12 h. Experimental protocols were opportunely approved by the Institutional Animal Care and Use Committee (protocol #008-16) and were performed according to national law #18.611 and international guidelines. All procedures were performed under Biosafety level II conditions.

Female mice were randomly distributed in three experimental groups: Infected (n=20), Infected + IVM (n=20) and Control (n=16). Experiments were conducted in three independent replicates.

### MHV-A59 preparation

MHV-A59 (ATCC^®^ VR-764™) viruses were expanded in murine L929 cells (ATCC^®^ CCL-1™) to reach a concentration of 1×10^7^ plaque forming unit (PFU)/mL. The virus-containing supernatants were stored at −80°C until further use.

### Infection and treatment

Before the infection, mice were weighed and bled from the submandibular vein for basal blood determinations. Mice were infected with 6,000 PFU of MHV-A59 diluted in 100 μL of sterile PBS administered by intraperitoneal route. Immediately after, mice from Infected + IVM group were treated with one single dose of 500 μg/kg of ivermectin (Ivomec 1%, Merial, France), diluted in 50 μL of PBS via s.c. The other two groups (Infected and Control) received 50 μL of PBS via s.c. Five days after infection/treatment, mice were weighed and 300 μL of blood were retrieved for plasma cytokines quantification, metabolic and hematological profile from submandibular vein. Mice were immediately euthanized by cervical dislocation to dissect liver and spleen for weight recording, histological and qPCR analysis. At necropsy, liver appearance was blindly scored (0 to 3) by an independent trained technician considering the main pathologic pattern of MHV infection (Macphee et al., 1985; Perlman, 1998). Briefly, gross hepatic lesions were identified as multifocal to coalescent whitish spots of less than 1mm diameter, defined as hepatic granulomas.

### Histological analysis

Immediately after necropsy, liver and spleen were fixed in 10% neutral buffered formalin (pH 7.4) for further processing. For evaluation, they were embedded in paraffin, sectioned at 4 μm and stained with hematoxylin-eosin (H&E), according to (Kyuwa et al., 2002). Specimens were whole examined under light microscope (Olympus BX41, Japan) at 10X in three randomly selected areas, or in the highest incidence areas of each specimen, by three different pathologists, to establish a histopathological score in each case, with a previously defined semi-quantitative microscopic grading centered in the identification of the typical histopathologic changes caused by MHV (characterized by the presence of hepatocellular necrotic areas and granulomatous inflammatory reaction), according to the following criteria: 0 = normal (no necrotic areas identified in the whole specimen); + = < 10 necrotic areas; ++ = 10-20 necrotic areas; +++ = > 20 necrotic areas.

### Hepatic viral load

After dissecting and trimming the whole liver, two samples (0.5×0.5cm, each) of the hepatic right lobe were retrieved for qPCR analysis. Samples were loaded in cryotubes with TRI Reagent® (Sigma-Aldrich, Saint-Louis, MO, US) and immediately plunged into liquid nitrogen until analysis. Total RNA was isolated according to the manufacturer’s instructions. cDNA was synthesized from 2μg total RNA, employing M-MLV Reverse Transcriptase (Thermo Fisher, Waltham, MA, USA) and random primers (Invitrogen, Carlsbad, CA, USA). Sample analysis was performed with a QuantStudio 3 Real-time PCR system (Thermo Fisher) using FastStart Universal SYBR Green Master (Rox) (Roche, Basel, CH). Primers employed sequences were MHV Forward primer (5’-3’): GGAACTTCTCGTTGGGCATTATACT and MHV Reverse primer (5’-3’): ACCACAAGATTATCATTTTCACAACATA. The reactions were performed according to the following settings: 95°C for 10 min, and 40 cycles of 95°C for 15 sec, followed by 60°C for 1 min. The quantification of viral loads was performed with a relative standard curve method.

### Blood biochemistry profile

Individual whole blood (100 μL) was analyzed for liver and kidney profile using the Pointcare V2 automatic device (Tianjin MNCHIP Technologies Co, China) at the beginning (preinfection determination) and at the end of the experiment (postinfection determination). Analyzed parameters included total proteins (TP), albumin (ALB), globulin (GLO), ALB/GLO ratio, total bilirubin (TBIL), alanine aminotransferase (ALT), aspartate aminotransferase (AST), gamma glutamiltranspeptidase (GGT), blood urea nitrogen (BUN), creatinine (CRE), BUN/CRE ratio and glucose (GLU).

### Hematological parameters

For hematologic analysis, aliquots of 20 μL of blood were collected into 0.5 mL microtubes containing EDTA potassium salts (W anticoagulant, Wiener lab, Rosario, Argentina) in a ratio of 1:10 (EDTA: blood) at pre and postinfection stages. All measurements were conducted within four hours after collection. Total white blood cells (WBC) count, differential WBC count and percentage, red blood cells (RBC) count, hemoglobin (HGC), hematocrit (HCT), and platelet (PLT) count, were evaluated using the auto hematology analyzer BC-5000Vet (Mindray Medical International Ltd., Shenzhen, China).

### Cytokines quantification and flow cytometry analysis

Bead-based multiplex assays were employed to quantify cytokines (LEGENDplex™ mouse Inflammation Panel, BioLegend Inc., San Diego, CA, USA) in plasma samples obtained from mice at pre and postinfection stages, according to manufacturer’s instructions. Briefly, blood samples with EDTA as anticoagulant were centrifuged for 10 min at 1000 x g, and plasma was recovered and stored at −20 °C until use. For the assay, 25 μL of 2-fold diluted plasma samples, diluted standards, and blanks were added into the corresponding tubes; 25 μL of pre-mixed beads and detection antibodies were added to all the tubes. Tubes were incubated for 2 h at room temperature with shaking. After this, and without washing, 25 μL of StreptAvidin-PhycoErythrin (SA-PE) conjugate was added, and the tubes were incubated for 30 min and finally washed and suspended in 200 μL of wash buffer. Data were acquired in a BD AccuriTM C6 (BD Biosciences, USA) flow cytometer. BD AccuriTM C6 software was used for data acquisition. Beads excitation was achieved using 488 and 640 nm lasers and emission was detected using 530/30 and 665/20 nm bandpass filters, respectively. For each analyte to be detected, 4,000 beads gated on a forward scatter (FSC) versus side scatter (SSC) dot plot were recorded. Data were processed with BioLegend LEGENDplex™ Data Analysis Software. Results represent the concentration expressed in pg/mL.

### Lymphocytes B and T analysis by flow cytometry

Lymphocytes surface markers were evaluated in peripheral blood samples (50 μL) anticoagulated with EDTA. Erythrocytes were removed by suspending cells in 1 mL of lysis buffer (155 mM NH4Cl, 12mM NaHCO3, 0.1mM EDTA, pH 7.4) for 10 min at room temperature. After washing in PBS containing 0.2% bovine serum albumin, nucleated cells were incubated on ice for 15 min with an antibody mixture. The following fluorophore-conjugated antibodies were used: Anti-CD4-FITC (#11004181, clone GK1.5) and anti-CD8-PE-Cy7 (#25008182, clone 53-6.7) from eBioscience™ (San Diego, CA, USA) and anti-CD19-PerCP-Cy™ 5.5 (#551001, clone ID3), from BD Pharmingen (San Diego, CA, USA). Flow cytometry analysis was performed using an Attune™ Nxt Acoustic Focusing Cytometer (Thermo Fisher) equipped with a 488 nm laser. Emissions were detected using 530/30, 695/40 and 780/60 nm bandpass filters, for FITC, PerCP-Cy5.5 and PE-Cy7, respectively. FlowJo™ software, version 10.6.1 (Tree star, Ashland, Oregon, USA) was used for data analysis. Unstained controls, single-color controls and fluorescence- minus-one controls were used to compensate and to establish baseline gate settings for each respective antibody combination.

Lymphocytes were gated based on their FSC and SSC dot plot profile, and FSC area vs FSC height dot plot was used to exclude doublets. B lymphocytes were defined as CD19-PerCP-Cy5.5 positive cells. For T lymphocyte analysis, a gate was placed on CD19-negative population and based on PE-Cy7 vs FITC dot plot, CD8-PE-Cy7 positive cells and CD4-FITC positive cells were defined as CD8+ and CD4+ lymphocytes, respectively. A minimum of 10,000 events in a single cell region were collected. Results were expressed as % of specific cell type from analyzed single cell population

### Statistical analysis

Statistical analysis was performed by using generalized linear mixed models (GLMM, InfoStat software (Di Rienzo et al., 2017), which included the treatments (three groups) and time (pre and postinfection) as fixed variables and the animals and replicates as random variables. Data were checked for normality and homogeneity of variance by histograms, q-q plots, and formal statistical tests as part of the univariate procedure. The type of variance-covariance structures was chosen depending on the magnitude of the Akaike information criterion (AIC) for models run under heterogeneous compound symmetry, unstructured, autoregressive, spatial power, and first-order ante-dependence. The model with the lowest AIC was chosen. Data are presented as mean ± SEM and the significance level were defined for a *p*-value of 0.05.

## Results

### Body and organ weight, macroscopic liver appearance

Body weight determined at the beginning and at the end of the experiment (*i.e*., pre and postinfection, respectively) was affected by the viral infection. While those animals from Control and Infected + IVM groups gained weight during the experimental period (p<0.05), those mice from the Infected group did not show statistical differences pre and postinfection (P=NS). Both liver and spleen from Infected and Infected + IVM groups were heavier than Control group at necropsy (p<0.05). Liver macroscopic appearance was impaired in the Infected group when compared to Infected + IVM or Control group (p<0.05). Results are shown in Figure 1 and 2.

**Figure 1:**
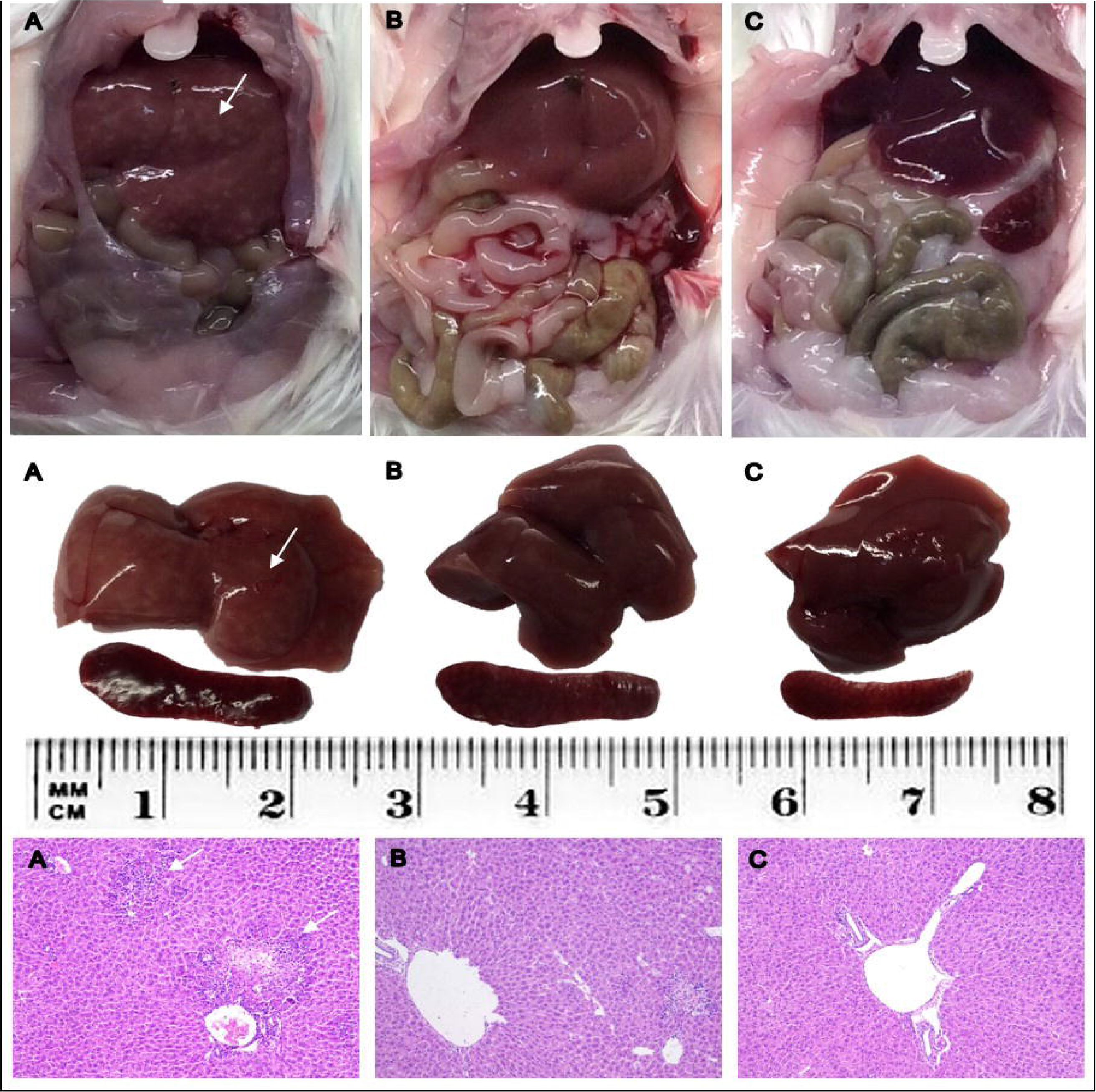
Representative liver and spleen from each group: A) Infected; B) Infected + IVM; C) Control. Upper panel: abdominal cavity at necropsy; middle panel: dissected liver and spleen; lower panel: HE histological sections of livers. White arrows indicate white spotted patterns in the liver from infected mice, and severe hepatocellular necrosis and lymphoplasmacytic inflammatory infiltration in histological images (A). IVM: ivermectin.

**Figure 2:**
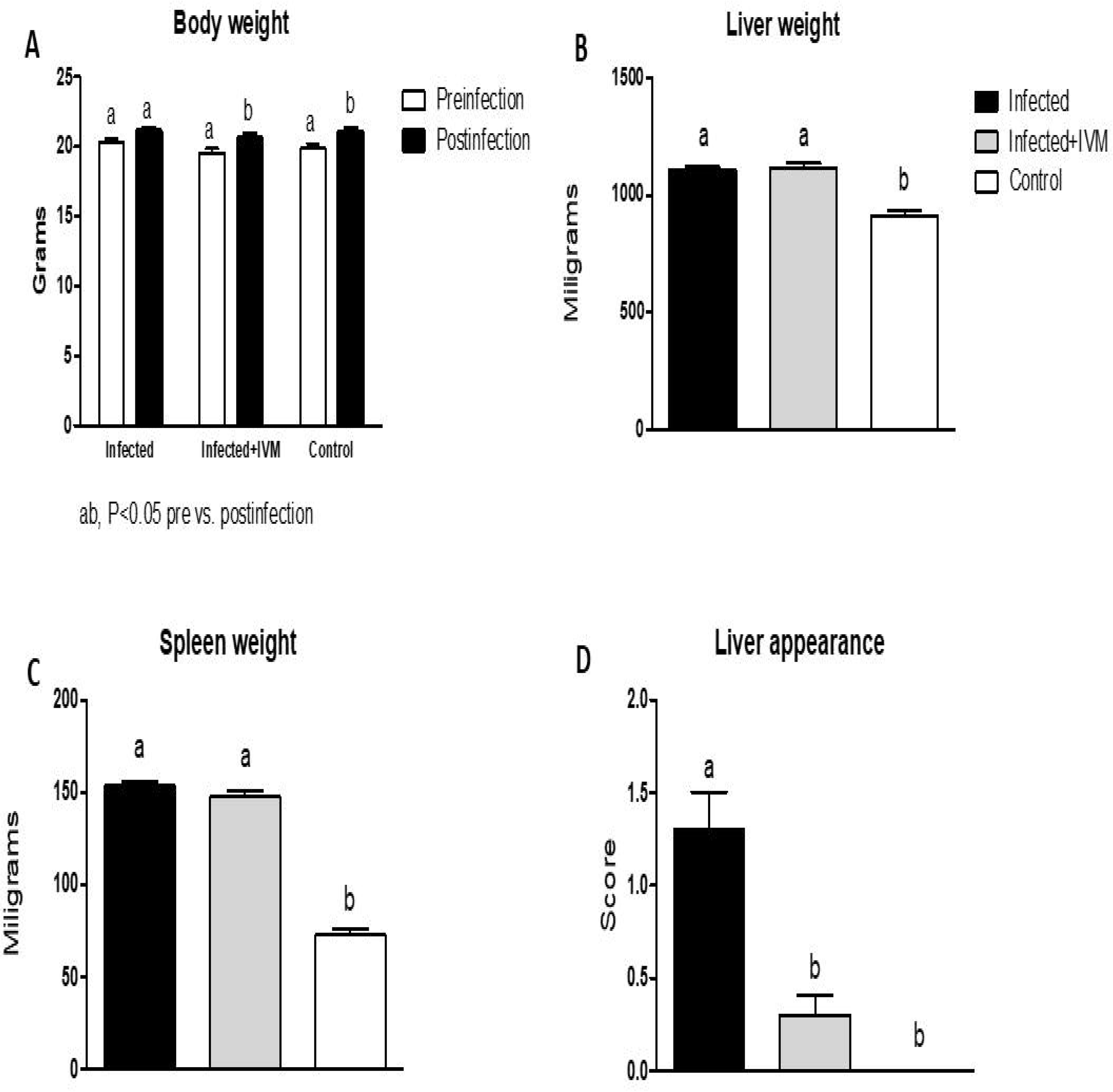
Body weight at the beginning and the end of the experiment (A), and organ weight and liver appearance at necropsy five days postinfection (B, C and D, respectively). Both liver and spleen of infected animals were heavier than control group (p<0.05). Liver macroscopic appearance was impaired in mice from the Infected group when compared with mice from the Infected + IVM or Control group (scale 0-3) (p<0.05) (Mean ± SD). Different letters indicate significant differences (p<0.05) between pre and post infection time for A, and significant differences (p<0.05) between indicated groups for B, C and D.

### Histological analysis

As observed, livers of mice from the Infected group were characterized by a severe hepatocellular necrosis, with high incidence of specimens (15/20) with more than 20 necrotic areas, surrounded by a severe lymphoplasmacytic inflammatory infiltration. The typical hepatocellular necrosis and inflammatory infiltration were present, but in all the cases of lesser grading in the Infected + IVM group (6/20). Mice from Control group did not show any hepatocellular or spleen lesions (0/16). Representative histological liver images are shown in Figure 1.

### Hepatic viral load

Results obtained from qPCR analysis showed a significantly higher viral load in the livers of Infected group vs. Infected + IVM or Control group (p<0.05), with a load of 52,158 ± 15,235 arbitrary units (AU) for the infected animals (Figure 3). Ivermectin treatment significantly lowered the viral load (23,192 ± 6,796 AU; p<0.05), which is in well accordance with other disease features observed.

**Figure 3:**
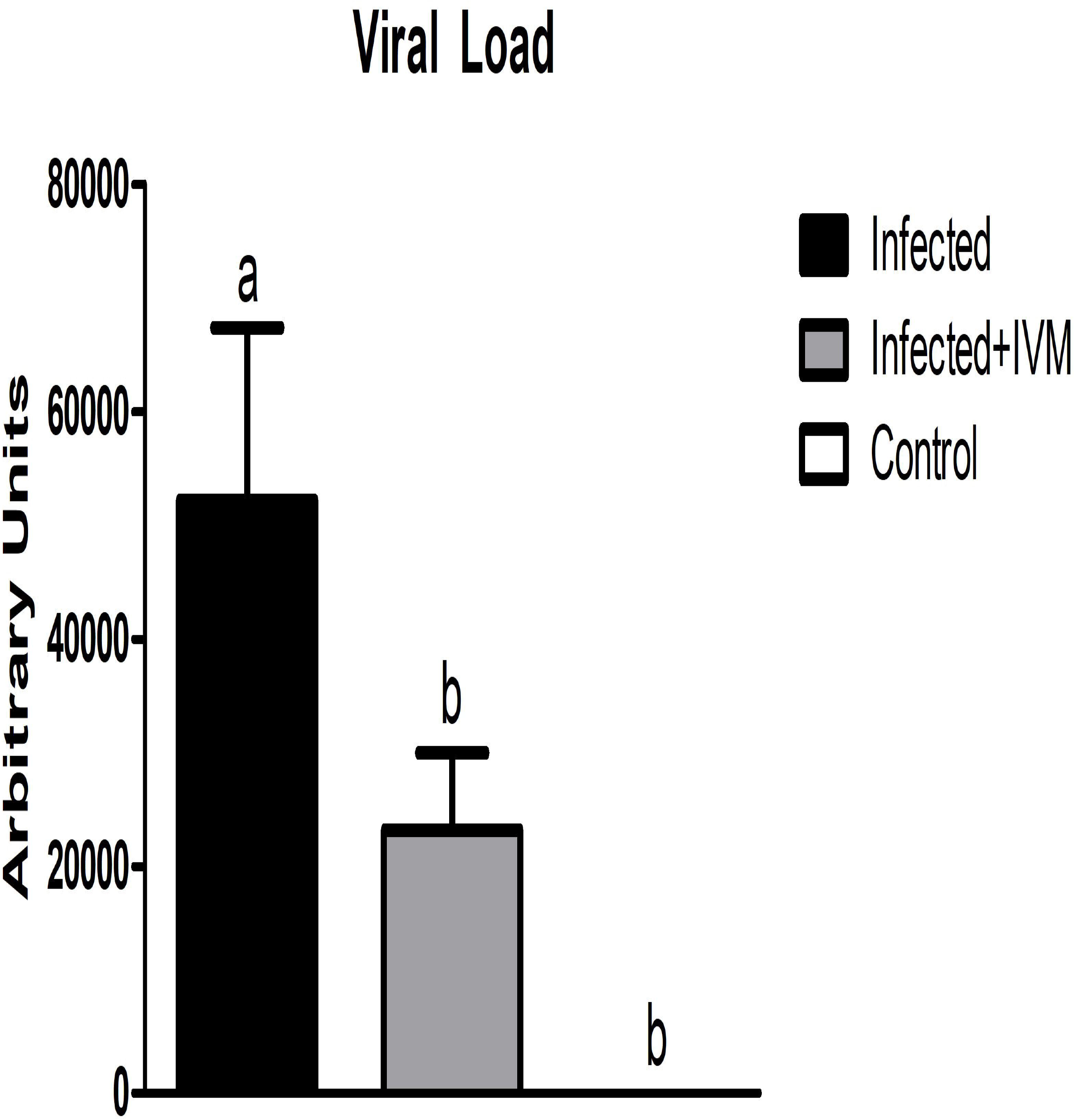
Hepatic viral load measured by qPCR in samples of liver from the three experimental groups. Viral load (expressed in arbitrary units) is higher in mice from the Infected group when compared with mice from Infected + IVM or Control group (p<0.05) (Mean ± SD). Different letters indicate significant differences (p<0.05) between groups.

### Blood biochemistry profile

To check liver and kidney health profile, several biochemical parameters (TP, GLO, ALB, A/G, TB, ALT, AST, GGT, BUN, CRE, BUN/CRE and GLU) were analyzed at the beginning (preinfection) and at the end of the experiment (postinfection) from complete blood. Total proteins decreased after infection in the Infected and Infected + IVM groups when compared with basal profile determinations (p<0.05) (Figure 4A). Albumin levels decreased in all experimental groups, being lower for Infected and Infected + IVM groups (p<0.05) (Figure 4B). On the contrary, globulin levels increased after infection for both infected groups, showing statistical differences when compared to Control group (p<0.05) (Figure 4C). The A/G ratio showed a decrease between basal and final stages for the three groups, again being lower for both infected groups (p<0.05) (Figure 4D). Total bilirubin did not show statistical differences among the groups for basal and final determinations (Figure 4E).

**Figure 4:**
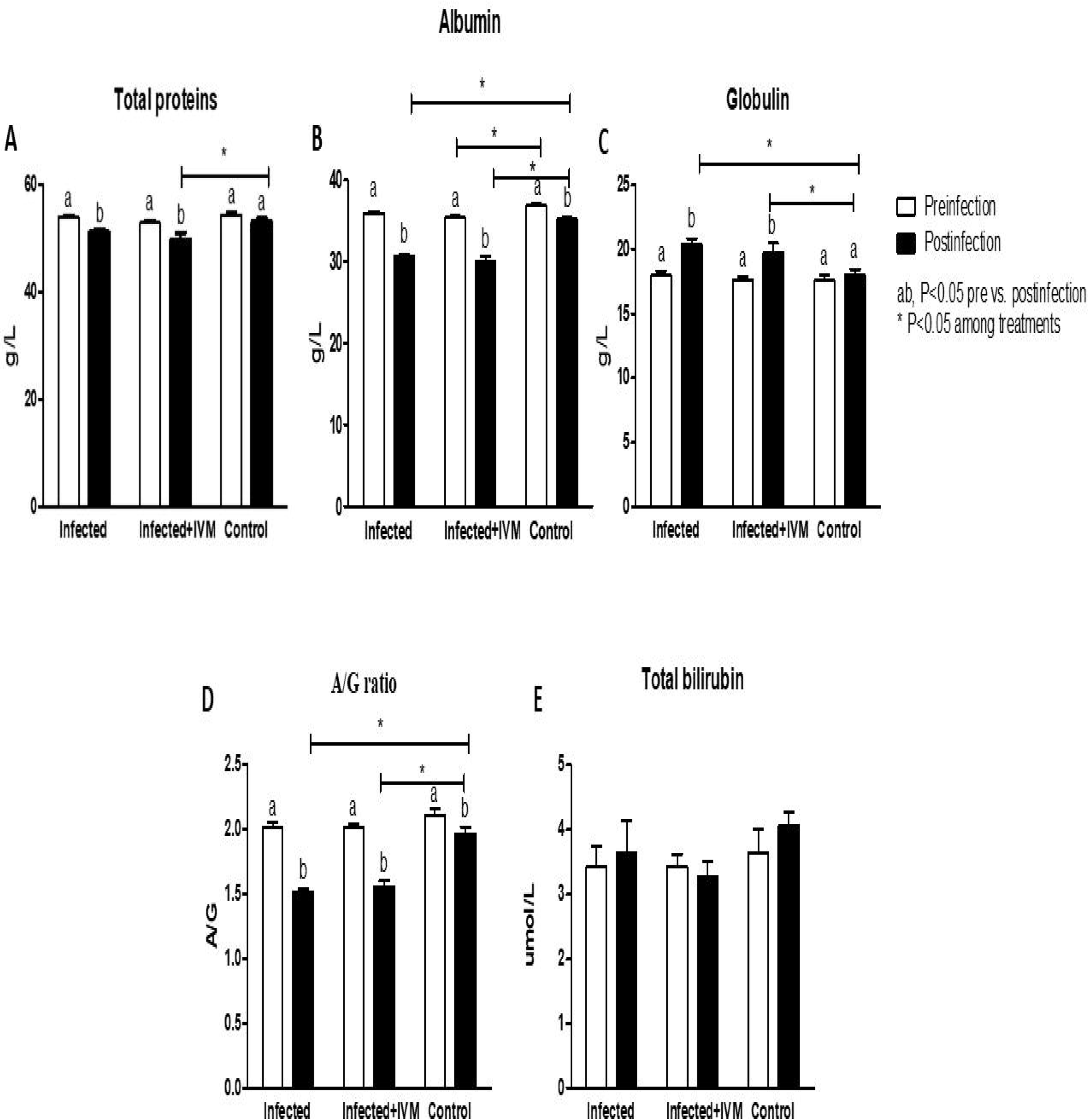
Protein levels measured in complete blood of the three experimental groups, before and after the infection with MHV A-59. A) Total proteins; B) Albumin; C) Globulin, D) Albumin/Globulin ratio and E) Total bilirubin (Mean ± SD). Different letters indicate significant differences (p<0.05) between pre and post infection time; asterisk (*) refers to significant differences (p<0.05) between indicated groups.

Hepatic transaminases such as ALT and AST showed an important increase in animals from the Infected group when compared with animals from the Infected + IVM or Control group (p<0.05) (Figure 5A and B). GGT levels were similar among groups and pre or postinfection (P=NS) (Figure 5C). BUN levels decreased after infection/treatment (p<0.05). CRE levels were lower for the animals of the Infected group at the end of the experiment when compared with animals from the Infected + IVM group (p<0.05). BUN/CRE ratio was similar among groups and blood determinations (P=NS), and GLU levels decreased in animals from the Infected group after infection when compared with Control group (p<0.05). Results are shown in Figure 6A, B, C and D.

**Figure 5:**
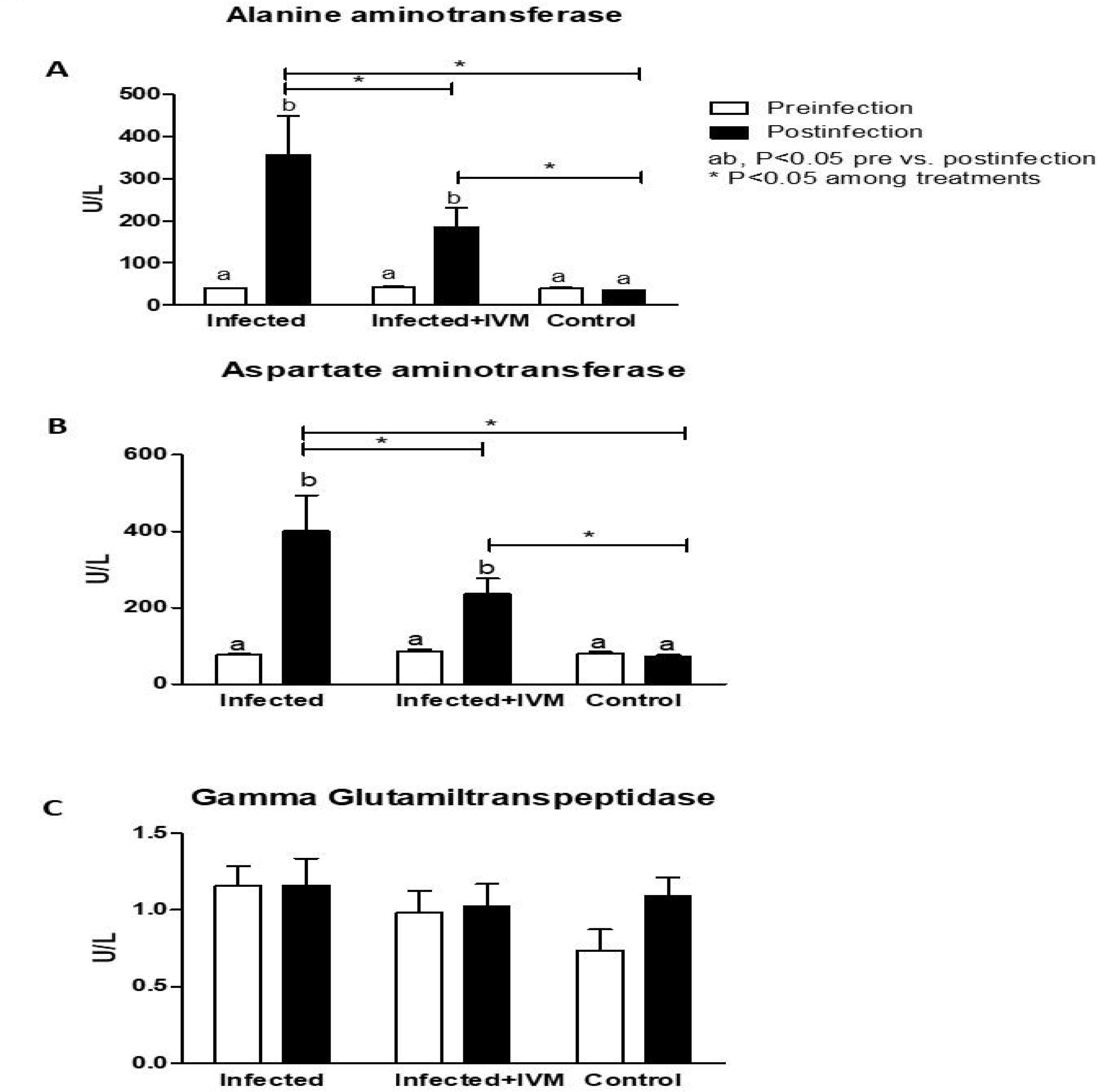
Hepatic enzymes levels obtained from blood of the three experimental groups, before and after virus infection. A) Alanine aminotransferase; B) Aspartate aminotransferase and C) Gamma Glutamiltranspeptidase (Mean ± SD). Different letters indicate significant differences (p<0.05) between pre and post infection time; asterisk (*) refers to significant differences (p<0.05) between indicated groups.

**Figure 6:**
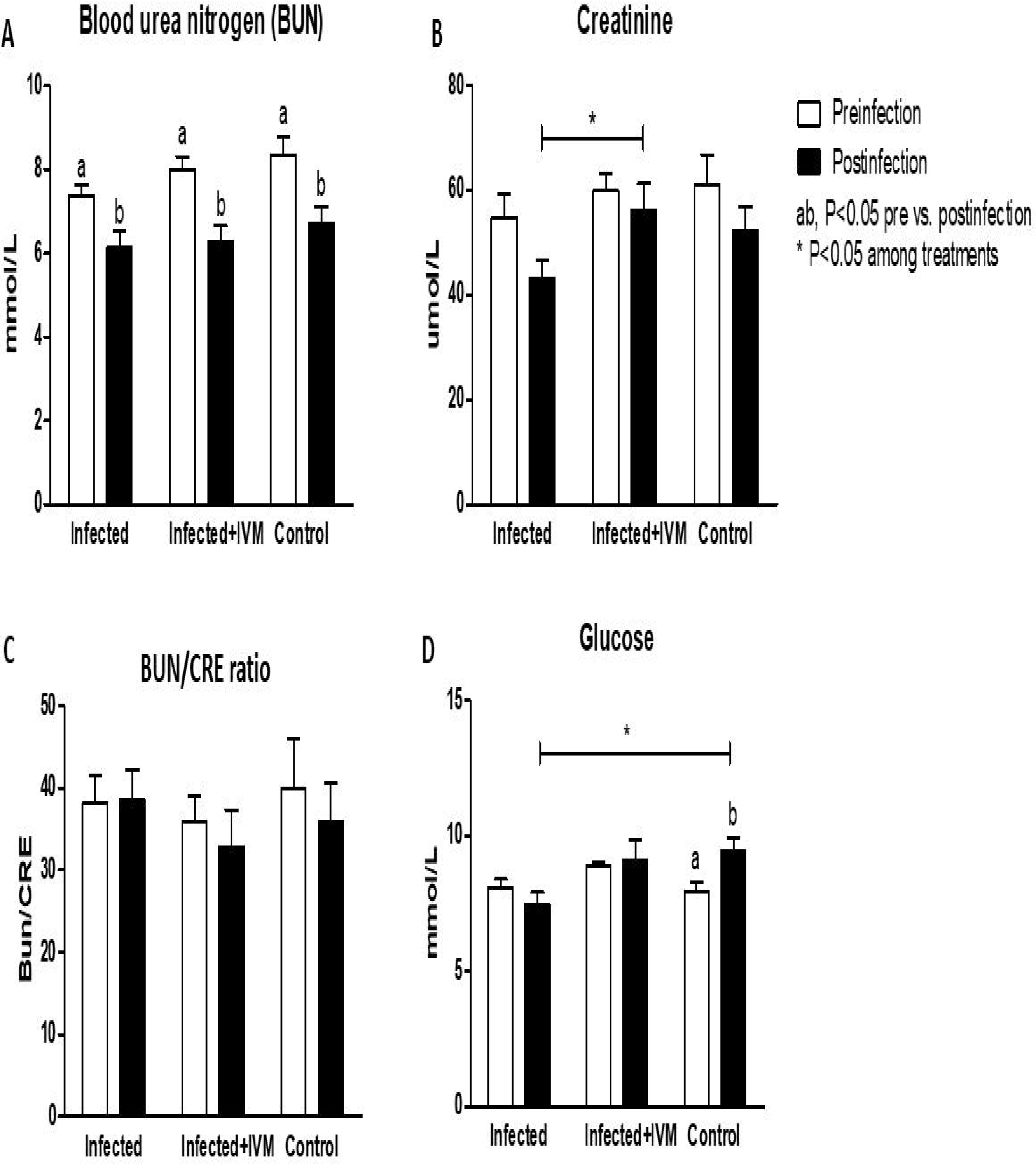
Blood urea nitrogen (BUN), creatinine (CRE), BUN/CRE ratio and glucose (GLU) levels were measured before and after infection. Results are shown in panel A, B, C and D, respectively (Mean ± SD). Different letters indicate significant differences (p<0.05) between pre and post infection time; asterisk (*) refers to significant differences (p<0.05) between indicated groups.

### Blood profile

Hematological parameters obtained from peripheral blood samples at pre and postinfection stages are summarized in Table 1. A significant decrease in the number of WBC was found in animals from both virus-infected groups (Figure 7A), whereas no changes in RBC, HGC, HCT, or platelets were found, compared with control or pre-infection values (Table 1). Absolute WBC and lymphocyte counts were increased in the control group, comparing pre and postinfection time points.

**Table 1:**
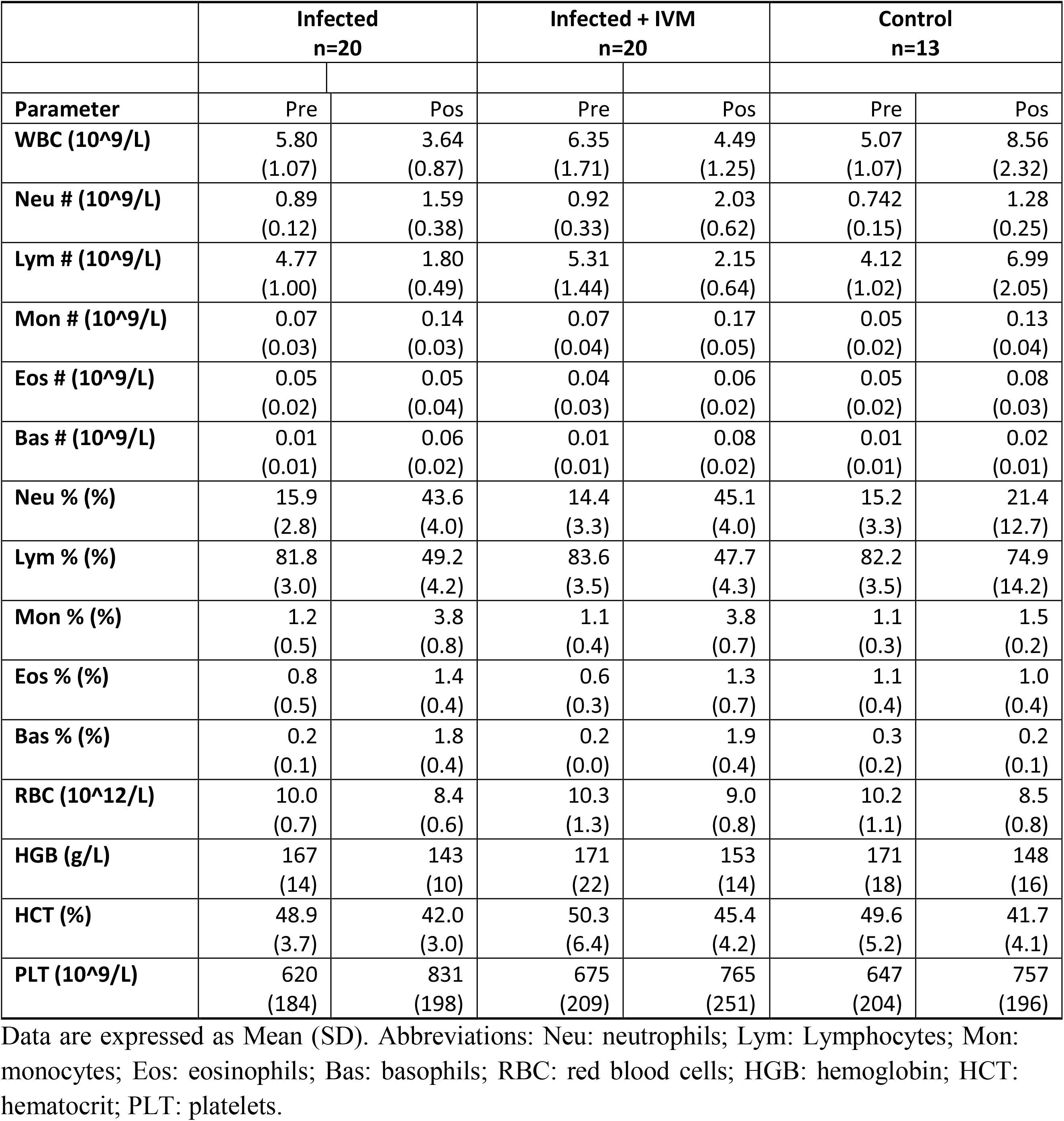
Hematological parameters from peripheral blood samples of the three experimental groups, measured at pre and postinfection time points.

|  | Infected<br>n=20 |  | Infected + IVM<br>n=20 |  | Control<br>n=13 |  |
| --- | --- | --- | --- | --- | --- | --- |
| Parameter | Pre | Pos | Pre | Pos | Pre | Pos |
| WBC (10<sup>9</sup>/L) | 5.80<br>(1.07) | 3.64<br>(0.87) | 6.35<br>(1.71) | 4.49<br>(1.25) | 5.07<br>(1.07) | 8.56<br>(2.32) |
| Neu # (10<sup>9</sup>/L) | 0.89<br>(0.12) | 1.59<br>(0.38) | 0.92<br>(0.33) | 2.03<br>(0.62) | 0.742<br>(0.15) | 1.28<br>(0.25) |
| Lym # (10<sup>9</sup>/L) | 4.77<br>(1.00) | 1.80<br>(0.49) | 5.31<br>(1.44) | 2.15<br>(0.64) | 4.12<br>(1.02) | 6.99<br>(2.05) |
| Mon # (10<sup>9</sup>/L) | 0.07<br>(0.03) | 0.14<br>(0.03) | 0.07<br>(0.04) | 0.17<br>(0.05) | 0.05<br>(0.02) | 0.13<br>(0.04) |
| Eos # (10<sup>9</sup>/L) | 0.05<br>(0.02) | 0.05<br>(0.04) | 0.04<br>(0.03) | 0.06<br>(0.02) | 0.05<br>(0.02) | 0.08<br>(0.03) |
| Bas # (10<sup>9</sup>/L) | 0.01<br>(0.01) | 0.06<br>(0.02) | 0.01<br>(0.01) | 0.08<br>(0.02) | 0.01<br>(0.01) | 0.02<br>(0.01) |
| Neu % (%) | 15.9<br>(2.8) | 43.6<br>(4.0) | 14.4<br>(3.3) | 45.1<br>(4.0) | 15.2<br>(3.3) | 21.4<br>(12.7) |
| Lym % (%) | 81.8<br>(3.0) | 49.2<br>(4.2) | 83.6<br>(3.5) | 47.7<br>(4.3) | 82.2<br>(3.5) | 74.9<br>(14.2) |
| Mon % (%) | 1.2<br>(0.5) | 3.8<br>(0.8) | 1.1<br>(0.4) | 3.8<br>(0.7) | 1.1<br>(0.3) | 1.5<br>(0.2) |
| Eos % (%) | 0.8<br>(0.5) | 1.4<br>(0.4) | 0.6<br>(0.3) | 1.3<br>(0.7) | 1.1<br>(0.4) | 1.0<br>(0.4) |
| Bas % (%) | 0.2<br>(0.1) | 1.8<br>(0.4) | 0.2<br>(0.0) | 1.9<br>(0.4) | 0.3<br>(0.2) | 0.2<br>(0.1) |
| RBC (10<sup>12</sup>/L) | 10.0<br>(0.7) | 8.4<br>(0.6) | 10.3<br>(1.3) | 9.0<br>(0.8) | 10.2<br>(1.1) | 8.5<br>(0.8) |
| HGB (g/L) | 167<br>(14) | 143<br>(10) | 171<br>(22) | 153<br>(14) | 171<br>(18) | 148<br>(16) |
| HCT (%) | 48.9<br>(3.7) | 42.0<br>(3.0) | 50.3<br>(6.4) | 45.4<br>(4.2) | 49.6<br>(5.2) | 41.7<br>(4.1) |
| PLT (10<sup>9</sup>/L) | 620<br>(184) | 831<br>(198) | 675<br>(209) | 765<br>(251) | 647<br>(204) | 757<br>(196) |

Data are expressed as Mean (SD). Abbreviations: Neu: neutrophils; Lym: Lymphocytes; Mon: monocytes; Eos: eosinophils; Bas: basophils; RBC: red blood cells; HGB: hemoglobin; HCT: hematocrit; PLT: platelets.

**Figure 7:**
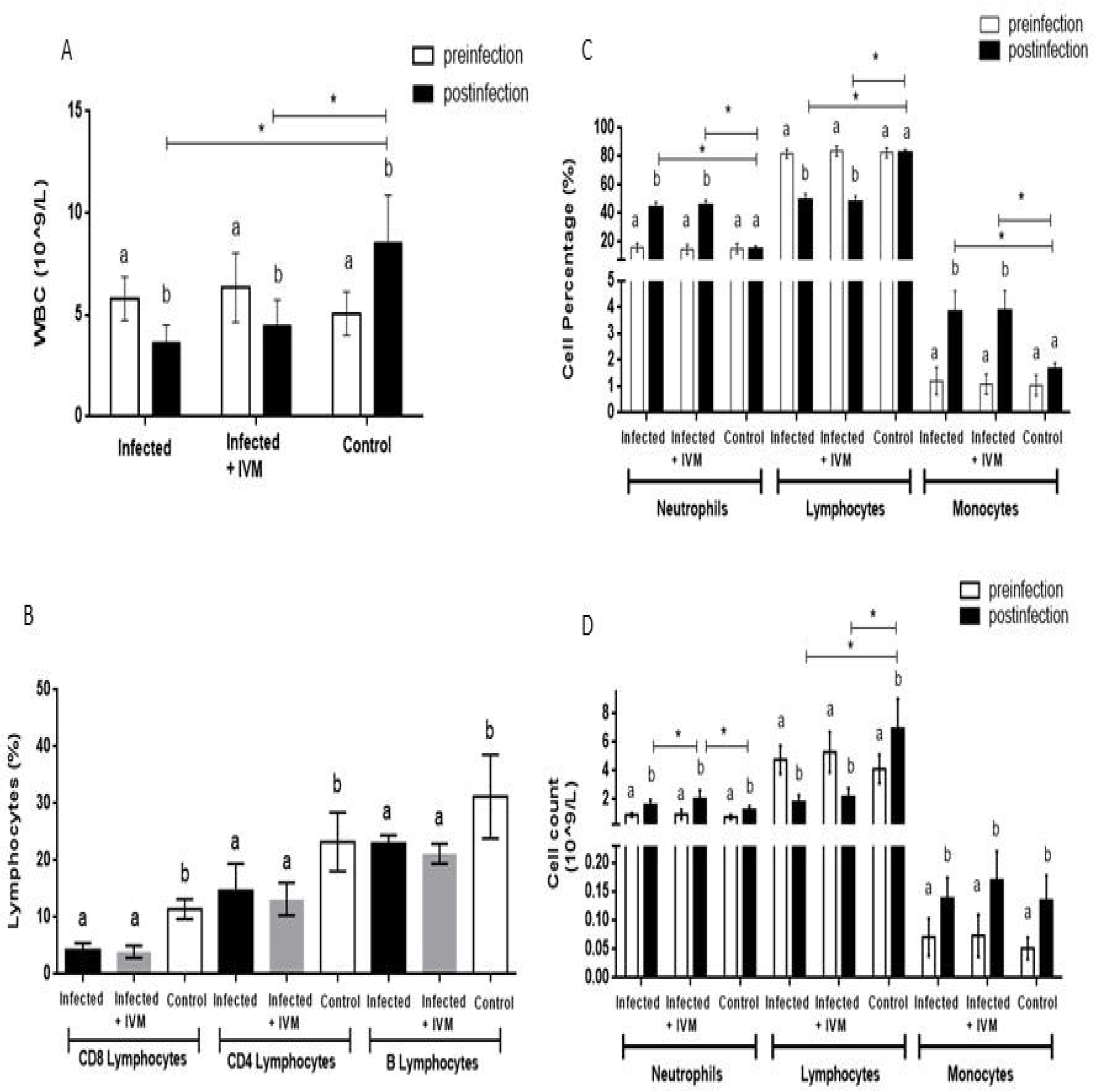
Evaluation of hematological parameters in mice. (A) White blood cells (WBC) count, (C, D) neutrophils, lymphocytes and monocytes counts and percentages, were determined in blood samples before and after infection for each experimental group (Infected, Infected+IVM, and Control) (Mean ± SD). Different letters indicate significant differences (p<0.05) between pre and postinfection time; asterisk (*) refers to significant differences (p<0.05) between indicated groups. (B) Lymphocytes staining for expression of surface markers for B and T cells at the end point of the experiment for all groups (Infected, Infected+IVM, and Control) (Mean ± SD) Different letters indicate statistically significant differences between groups (p<0.05).

Regarding WBC differential, the percentage of neutrophils and monocytes significantly increased in animals from both infected groups. In contrast, the percentage and counts of lymphocytes were the only WBC parameters significantly decreased in animals from infected groups, regardless of ivermectin treatment, when compared with the control or pre-infection values (Figure 7C). Moreover, animals from the infected group that received ivermectin treatment showed an increase in the number of neutrophils compared with animals from the infected group (Figure 7D).

To further characterize the reduction of lymphocyte population observed in animals from the infected groups, B and T lymphocytes were analyzed by the detection of specific cell surface markers: CD19 (B lymphocytes) or CD8 and CD4 (T lymphocytes), at the endpoint of the experiment. Results showed that both B and T lymphocytes percentages were reduced in mice from virus-infected groups, compared to control group (Figure 7B), being the CD8+ cells the subpopulation with the highest reduction (64 % and 66% of depletion for Infected and Infected + IVM groups, respectively).

### Proinflammatory cytokines

Cytokines obtained from plasma samples at the endpoint of the experiment (five days after the viral inoculation) were measured in the three groups. From the panel of 13 inflammatory related cytokines, only IFNɣ and MCP-1 were significantly increased in both infected groups (p<0.05), compared with the control values, regardless ivermectin treatment. On the other hand, TNFα values, which were increased in mice from the Infected group, were reduced in the animals that received ivermectin treatment, the latter being not statistically different from the control mice (Figure 8).

**Figure 8:**
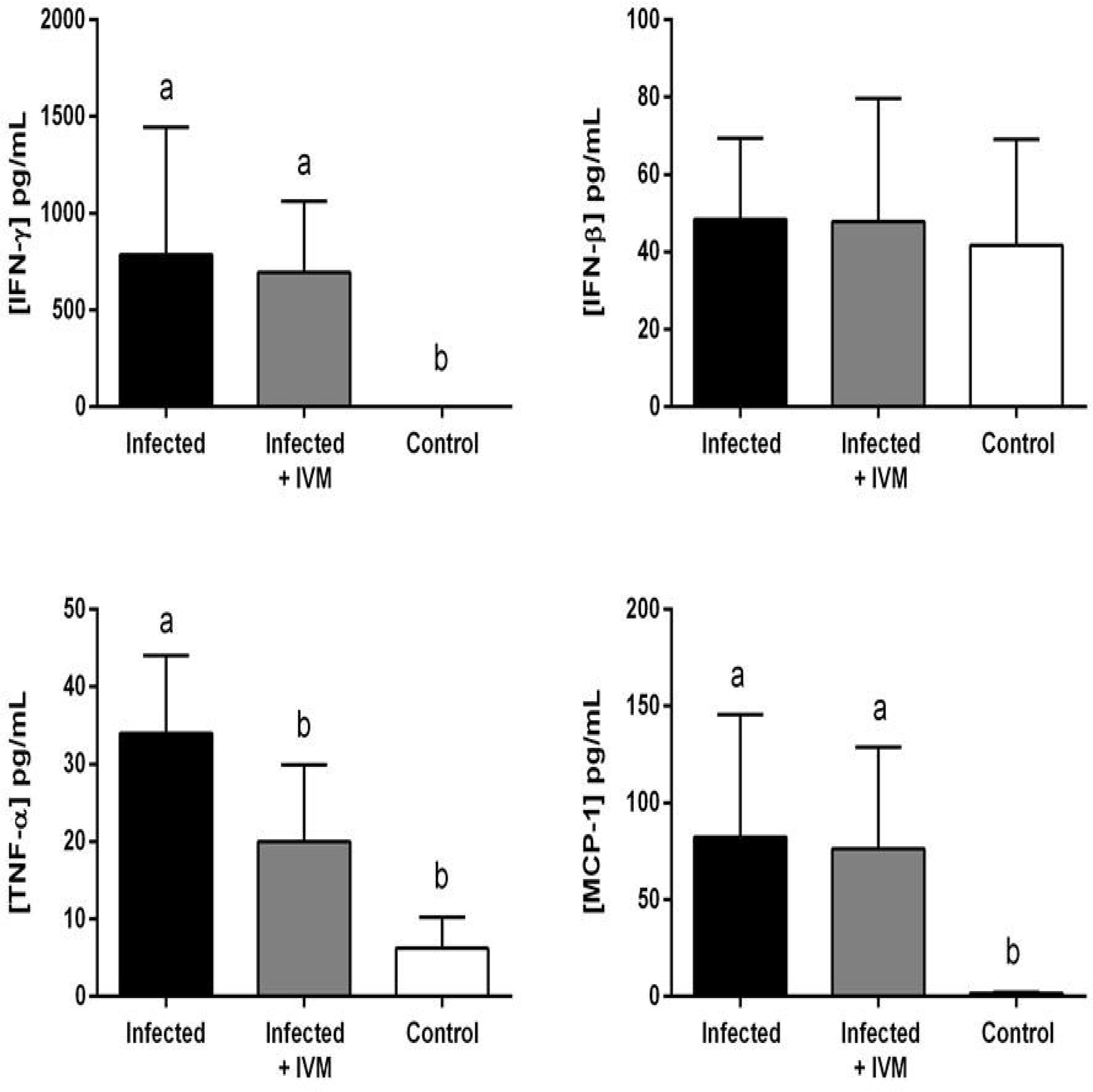
Detection of plasma cytokines. Murine plasma was obtained 5 days post infection and cytokine concentration was determined by multiplex bead array. (Mean ± SD). p<0.05. Different letters indicate statistically significant differences.

No statistically significant differences were observed for other analyzed cytokines (IL-1b, IL-1a, IL-23, IL-12p70, IL-10, or IFNβ) and for other cases (IL-6, IL-27, IL-17A and GM-CSF) proteins could not even be detected as their values were below the detection limit of the assay.

## Discussion

This study proposes a mouse experimental model for *in vivo* evaluation of pharmacological therapies against coronavirus diseases. It is well known that preclinical animal models are of utmost relevance when developing new therapies or vaccines that will be applied in humans. The need to develop animal models to study SARS-CoV2 has been recently proposed by many researchers (Johansen et al., 2020). Our study is based in the already tested *in vitro* reports of the use of ivermectin against several other RNA and DNA human and animal viruses (Heidary and Gharebaghi, 2020), such as influenza A virus, West Nile virus, Venezuelan equine encephalitis virus, Zika, chikungunya, Newcastle disease, porcine reproductive and respiratory syndrome virus, HIV-1, dengue virus, yellow fever and tick-borne encephalitis virus, pseudorabies, porcine circovirus, parvovirus and bovine herpesvirus. However, most of these studies reported only *in vitro* results and the information of the effect of this drug used in *in vivo* models is scarce. Regarding the recent appearance of SARS-CoV2, although several ongoing studies are being conducted, no information has been published yet on the *in vivo* effect of ivermectin administration on infected individuals with this kind of virus.

In our model, mice infected with MHV and immediately treated with ivermectin showed a lower hepatic viral load five days after infection, and a better general health status when compared with infected animals with no ivermectin treatment. At the moment of the necropsy and histological analysis, the liver of infected and untreated mice showed the worst appearance, with several animals with severe hepatocellular necrosis and lymphoplasmacytic inflammatory infiltration. Treated group showed lesser grading of anomalies, although liver and spleen weight were heavier for both infected groups when compared with the control group. The organ weight increase following the infection is indicative of the immune reaction (Robinson et al., 2016), something that cannot be evaluated in an *in vitro* model. These findings are commonly found in MHV infected mice (Barthold, 1997) and the generalized immune reaction can be confirmed with the cytokine’s levels found in our study in both infected groups. Viral load was significantly lower in those infected animals that received ivermectin, probably due to an impairment in virus entrance to the cell since this drug has been shown to inhibit nuclear import to the host cell (Kosyna et al., 2015).

The liver and kidney serum biochemical outcomes showed a clear impairment of metabolic profile mainly due to liver damage. Both groups of infected mice showed hypoalbuminemia and hyperglobulinemia, with a decrease in A/G ratio when compared with the control group. Variation in both proteins are indicative of hepatic damage (Carvalho and Machado, 2018). Serum concentration of transaminases such as AST and ALT, which are also indicative of liver function (Smith et al., 2013), were significantly higher in those virus infected mice that did not receive ivermectin, and associated with the rest of the studied variables suggest liver injury. A considerable decrease in serum creatinine levels was found in infected mice, again representing a major liver damage in sick animals. Glucose levels also showed a significant decrease in infected mice, probably due to the fasting of animals related to an impairment in general health status. All in all, this metabolic profile shows a major liver damage mostly in infected animals, in concordance with the rest of data analyzed during the study. The treatment with ivermectin was effective to reduce the effect of the virus infection, encouraging proposing novel therapies against coronavirus diseases.

The most relevant hematological findings were an increase in neutrophils and monocytes percentages and a reduction in WBC and lymphocytes (B and T) in both infected groups, regardless of ivermectin treatment. Neutrophilia and lymphopenia have been well documented in viral respiratory infection diseases in mouse models and humans (Camp and Jonsson, 2017; Feng et al., 2015; Preusse et al., 2015). The increase in the percentage of neutrophils in the virus-infected groups could be associated with the acute-phase viral infection. On the other hand, the reduction in lymphocytes might be due to migration/retention of these cells in the liver and/or lymphoid tissue. The rapid development of lymphopenia was also observed in COVID-19 patients with adverse outcomes, whereby CD4+ T-cells are more severely reduced than CD8+ T-cells (Chen and Subbarao, 2007; Guan et al., 2020). Neutrophils count was the only hematological parameter that differed among the virus-infected animals, being higher in mice from the ivermectin treated group. Nevertheless, this difference did not impact the WBC differential values. Moreover, in both infected groups, neutrophils counts were increased compared with the corresponding preinfection time point, and ivermectin treatment alone did not differ from control values (data not shown). Taking hematological data together, it seems that differences observed between groups would be related to the viral infection itself, rather than to an ivermectin effect. Studies about immunomodulatory effects of ivermectin are variable (Sajid et al., 2006), making it difficult to clearly define its function. On this regard, the *in vivo* mouse model of MHV infection would not support a modulatory action of ivermectin on the immune response. On the other hand, these results are in accordance with various reports demonstrating that the broad-spectrum antiviral potential of ivermectin against several RNA viruses is due to its ability to specifically bind to and destabilize the importin α/β heterodimer, thereby preventing importin α/β from binding to the viral protein, which in turn blocks the nuclear trafficking of viral proteins (Jans and Wagstaff, 2020; Sharun et al., 2020; Caly et al., 2020).

Regarding cytokines analysis, only IFN-ɣ and MCP-1 were increased in mice from the viral-infected groups, compared to mice from the Control group. These increases are in line with the general immune response associated with a viral infection. On the other hand, ivermectin treatment seemed not to exert a significant effect in the modulation of most of the inflammatory cytokines. An exception was TNF-α, whose value was significantly reduced in the ivermectin treated animals when compared with mice from the infected group. It has been reported that ivermectin can exert anti-inflammatory effects in *in vitro* cell models by downregulating NF-kB signaling pathways and regulating TNF-α, IL-1β and IL-10 (Ci et al., 2009), and in *in vivo* models by decreasing the production of TNF-α, IL-1ß and IL-6 (Zhang et al., 2008).

In the present work, neither IL-1β nor IL-10 or IL-6 were modulated by ivermectin. It is possible that differences regarding the experimental model, the route of infection and the time window of the measurements could account for these discrepancies, since in a living organism the immune response is influenced by more than one cellular component of the immunological system. Moreover, the similar hematological profiles of both infected groups suggest that the main antiviral effect of the ivermectin would not be through immunomodulatory actions.

## Conclusion

This study demonstrates that ivermectin administration reduces MHV liver viral load in infected mice, enhancing general health status. This preclinical model can be suitable to further study the effect of ivermectin in coronavirus infection, as a possible murine surrogate model, helping to find available treatments for COVID-19 and other coronavirus-related diseases.

## Acknowledgements

Experiments were carried out with genuine funds from *Institut Pasteur de Montevideo* and *FOCEM (MERCOSUR Structural Convergence Fund), COF 03/11*. MS, JMV, MH, MB and MC are members of *Sistema Nacional de Investigadores* (SNI).

## References

Barthold, S.W., 1997. Mouse Hepatitis Virus Infection, Liver, Mouse, in: Jones, T.C., Popp, J.A., Mohr, U. (Eds.), Digestive System. Springer Berlin Heidelberg, Berlin, Heidelberg, pp. 179–184. https://doi.org/10.1007/978-3-662-25996-2_25

Caly, L., Druce, J.D., Catton, M.G., Jans, D.A., Wagstaff, K.M., 2020. The FDA-approved drug ivermectin inhibits the replication of SARS-CoV-2 in vitro. Antiviral Research 178, 104787. https://doi.org/10.1016/j.antiviral.2020.104787

Camp, J.V., Jonsson, C.B., 2017. A Role for Neutrophils in Viral Respiratory Disease. Front Immunol 8. https://doi.org/10.3389/fimmu.2017.00550 Vol. 17. Issue 4.

Carvalho, J and Machado M., 2018. New Insights About Albumin and Liver Disease, Annals of Hepatology, 547–560. DOI: 10.5604/01.3001.0012.0916

Ci X, Li H, Yu Q, Zhang X, Yu L, Chen N, Song Y, Deng X. 2009. Avermectin exerts anti-inflammatory effect by downregulating the nuclear transcription factor kappa-B and mitogen-activated protein kinase activation pathway. Fundam Clin Pharmacol. Aug;23(4):449–55. doi: 10.1111/j.1472-8206.2009.00684.x. Epub 2009 May 6. PMID: 19453757.

Crump, A., Ōmura, S., 2011. Ivermectin, ‘Wonder drug’ from Japan: the human use perspective. Proc Jpn Acad Ser B Phys Biol Sci 87, 13–28. https://doi.org/10.2183/pjab.87.13

Chen J, Subbarao K. 2007. The Immunobiology of SARS*. Annu Rev Immunol. 25:443–72. doi: 10.1146/annurev.immunol.25.022106.141706. PMID: 17243893.

Di Rienzo, J., Macchiavelli, R., Casanoves, F., 2017. Modelos lineales generalizados mixtos: aplicaciones en InfoStat.

Fan, X., Cao, D., Kong, L., Zhang, X., 2020. Cryo-EM analysis of the post-fusion structure of the SARS-CoV spike glycoprotein. Nature Communications 11, 3618. https://doi.org/10.1038/s41467-020-17371-6

Feng, Y., Hu, L., Lu, S., Chen, Q., Zheng, Y., Zeng, D., Zhang, J., Zhang, A., Chen, L., Hu, Y., Zhang, Z., 2015. Molecular pathology analyses of two fatal human infections of avian influenza A(H7N9) virus. J Clin Pathol 68, 57–63. https://doi.org/10.1136/jclinpath-2014-202441

Guan WJ, Ni ZY, Hu Y, Liang WH, Ou CQ, He JX, Liu L, Shan H, Lei CL, Hui DSC, Du B, Li LJ, Zeng G, Yuen KY, Chen RC, Tang CL, Wang T, Chen PY, Xiang J, Li SY, Wang JL, Liang ZJ, Peng YX, Wei L, Liu Y, Hu YH, Peng P, Wang JM, Liu JY, Chen Z, Li G, Zheng ZJ, Qiu SQ, Luo J, Ye CJ, Zhu SY, Zhong NS; China Medical Treatment Expert Group for Covid-19. Clinical Characteristics of Coronavirus Disease 2019 in China. N Engl J Med. 2020 Apr 30;382(18):1708–1720. doi: 10.1056/NEJMoa2002032. Epub 2020 Feb 28. PMID: 32109013; PMCID: PMC7092819.

Heidary, F., Gharebaghi, R., 2020. Ivermectin: a systematic review from antiviral effects to COVID-19 complementary regimen. The Journal of Antibiotics 73, 593–602. https://doi.org/10.1038/s41429-020-0336-z

Jans, D.A., Wagstaff, K.M., 2020. Ivermectin as a Broad-Spectrum Host-Directed Antiviral: The Real Deal? Cells 9, 2100. https://doi.org/10.3390/cells9092100

Johansen, M.D., Irving, A., Montagutelli, X., Tate, M.D., Rudloff, I., Nold, M.F., Hansbro, N.G., Kim, R.Y., Donovan, C., Liu, G., Faiz, A., Short, K.R., Lyons, J.G., McCaughan, G.W., Gorrell, M.D., Cole, A., Moreno, C., Couteur, D., Hesselson, D., Triccas, J., Neely, G.G., Gamble, J.R., Simpson, S.J., Saunders, B.M., Oliver, B.G., Britton, W.J., Wark, P.A., Nold-Petry, C.A., Hansbro, P.M., 2020. Animal and translational models of SARS-CoV-2 infection and COVID-19. Mucosal Immunology 13, 877–891. https://doi.org/10.1038/s41385-020-00340-z

Körner, R.W., Majjouti, M., Alcazar, M.A.A., Mahabir, E., 2020. Of Mice and Men: The Coronavirus MHV and Mouse Models as a Translational Approach to Understand SARS-CoV-2. Viruses 12. https://doi.org/10.3390/v12080880

Kosyna, F., Nagel, M., Kluxen, L., Kraushaar, K., Depping, R., 2015. The importin alpha/beta-specific inhibitor Ivermectin affects HIF-dependent hypoxia response pathways. Biol. Chem. 396, 1357–1367.

Kyuwa, S., Shibata, S., Tagawa, Y., Iwakura, Y., Machii, K., Urano, T., 2002. Acute hepatic failure in IFN-γ-deficient BALB/c mice after murine coronavirus infection. Virus Res 83, 169–177. https://doi.org/10.1016/S0168-1702(01)00432-4

Macphee, P.J., Dindzans, V.J., Fung, L., Levy, G.A., 1985. Acute and chronic changes in the microcirculation of the liver in inbred strains of mice following infection with mouse hepatitis virus type 3. Hepatology 5, 649–660. https://doi.org/10.1002/hep.1840050422

Perlman, S., 1998. Pathogenesis of coronavirus-induced infections. Review of pathological and immunological aspects. Adv Exp Med Biol 440, 503–513.

Preusse, M., Schughart, K., Wilk, E., Klawonn, F., Pessler, F., 2015. Hematological parameters in the early phase of influenza A virus infection in differentially susceptible inbred mouse strains. BMC Research Notes 8, 225. https://doi.org/10.1186/s13104-015-1195-8

Robinson MW, Harmon C, O’Farrelly C. Liver immunology and its role in inflammation and homeostasis. Cell Mol Immunol. 2016 May;13(3):267–76. doi: 10.1038/cmi.2016.3. Epub 2016 Apr 11. PMID: 27063467; PMCID: PMC4856809.

Sajid, M.S., Iqbal, Z., Muhammad, G., Iqbal, M.U., 2006. Immunomodulatory effect of various anti-parasitics: a review. Parasitology 132, 301–313. https://doi.org/10.1017/S0031182005009108

Sharun, K., Dhama, K., Patel, S.K., Pathak, M., Tiwari, R., Singh, B.R., Sah, R., Bonilla-Aldana, D.K., Rodriguez-Morales, A.J., Leblebicioglu, H., 2020. Ivermectin, a new candidate therapeutic against SARS-CoV-2/COVID-19. Annals of Clinical Microbiology and Antimicrobials 19, 23. https://doi.org/10.1186/s12941-020-00368-w

Smith, G., Walter, G., Walker, R. 2013. Clinical Pathology in Non-Clinical Toxicology Testing. In: Haschek and Rousseaux’s Handbook of Toxicologic Pathology (Third Edition). Academic Press, Editor(s): Wanda M. Haschek, Colin G. Rousseaux, Matthew A. Wallig, Pages 565–594, ISBN 9780124157590, https://doi.org/10.1016/B978-0-12-415759-0.00018-2.

Timani, K.A., Liao, Q., Ye, Linbai, Zeng, Y., Liu, J., Zheng, Y., Ye, Li, Yang, X., Lingbao, K., Gao, J., Zhu, Y., 2005. Nuclear/nucleolar localization properties of C-terminal nucleocapsid protein of SARS coronavirus. Virus Res 114, 23–34. https://doi.org/10.1016/j.virusres.2005.05.007

Wagstaff, K.M., Sivakumaran, H., Heaton, S.M., Harrich, D., Jans, D.A., 2012. Ivermectin is a specific inhibitor of importin α/β-mediated nuclear import able to inhibit replication of HIV-1 and dengue virus. Biochem J 443, 851–856. https://doi.org/10.1042/BJ20120150

Weiss, S.R., Leibowitz, J.L., 2011. Chapter 4 - Coronavirus Pathogenesis, in: Maramorosch, K., Shatkin, A.J., Murphy, F.A. (Eds.), Advances in Virus Research. Academic Press, pp. 85–164. https://doi.org/10.1016/B978-0-12-385885-6.00009-2

Wulan, W.N., Heydet, D., Walker, E.J., Gahan, M.E., Ghildyal, R., 2015. Nucleocytoplasmic transport of nucleocapsid proteins of enveloped RNA viruses. Front. Microbiol. 6. https://doi.org/10.3389/fmicb.2015.00553

Zhang X, Song Y, Ci X, An N, Ju Y, Li H, Wang X, Han C, Cui J, Deng X. 2008. Ivermectin inhibits LPS-induced production of inflammatory cytokines and improves LPS-induced survival in mice. Inflamm Res. Nov;57(11):524–9. doi: 10.1007/s00011-008-8007-8. PMID: 19109745.

